# Dorsal hippocampus mediates light-tone associations in male mice

**DOI:** 10.1101/2025.01.07.631667

**Authors:** Julia S. Pinho, Carla Ramon-Duaso, Irene Manzanares-Sierra, Arnau Busquets-García

**Author notes:** Corresponding author: Arnau Busquets-Garcia, PhD, Cell-Type Mechanisms in Normal and Pathological Behavior Research Group, Neuroscience Research Program, Hospital del Mar Research Institute. PRBB, 88 Dr. Aiguader street Barcelona 08003 SPAIN.

## Abstract

Daily choices are often influenced by environmental cues that are not directly associated with reinforcers. This phenomenon, known as higher-order conditioning, can be studied using sensory preconditioning tasks in rodents. This behavioral paradigm involves the repeated pairing of two innocuous stimuli, such as a light and a tone, followed by a devaluation phase in which one stimulus is associated with an unconditioned stimulus, such as a mild foot-shock. The result is a conditioned response (e.g., freezing) to both the conditioned stimulus (direct learning) and the non-conditioned stimulus (mediated learning). In our study, we successfully established a light-tone sensory preconditioning task specifically in male mice, as sex differences were observed in both control experimental groups and in sensory preconditioning responses. We employed in vivo, freely moving fiber photometry to monitor neural activity in the dorsal and ventral subregions of the hippocampus in male mice during the formation of associations between innocuous stimuli and reinforcers. Additionally, we combined our sensory preconditioning task with chemogenetic approaches to investigate the roles of these hippocampal subregions in sensory preconditioning. Our results indicate that dorsal, but not ventral, CaMKII-positive neurons are involved in encoding innocuous stimuli during the preconditioning phase. Overall, we developed a novel light-tone sensory preconditioning protocol in male mice, enabling the detection of sex differences and furthering our understanding of how specific hippocampal subregions and cell types regulate complex cognitive processes.

## Introduction

The brain mechanisms underlying associative learning have traditionally been elucidated through classical conditioning paradigms involving salient stimuli as reinforcers—such as pairing an electric foot-shock with innocuous stimuli (e.g., a light, tone, or odor). However, these paradigms are not sufficient to fully capture the complexity of animal behavior, as daily choices are not always dictated by stimuli that have been directly associated with a potent reinforcer. In both humans and animals, responses often occur to cues that were never explicitly conditioned but had previously been associated with other stimuli linked to specific aversive or rewarding outcomes^1–5^.

These higher-order conditioning processes can be studied in laboratory settings using sensory preconditioning procedures^2,6–11^. These tasks involve incidental associations between neutral stimuli (e.g., odors, tastes, lights, or tones) during a preconditioning phase, followed by classical conditioning of one of these stimuli with an aversive or appetitive unconditioned reinforcer^2,6–11^. As a result, subjects exhibit aversion or preference towards the stimulus that was never directly paired with the reinforcer, enabling the assessment of mediated learning^2,6–11^. While the brain circuits underlying classical associative memory between neutral cues and reinforcers have been extensively studied, the mechanisms responsible for incidental associations between innocuous sensory stimuli—and their behavioral consequences—remain much less understood.

The hippocampus and other cortical regions (e.g., the perirhinal, retrosplenial, and orbitofrontal cortices) have been implicated in higher-order conditioned responses^6,8,10–14^. Sensory preconditioning tasks using various sensory modalities have shown that the involvement of different brain regions may depend on the behavioral phase being studied (i.e., preconditioning or testing) or the modality of the sensory cues (visual, gustatory, olfactory, or auditory). However, it is still unknown whether there are common brain regions where different types of stimuli are integrated, regardless of the behavioral phase or the modality. The hippocampus has been proposed to play a central role in multiple behavioral phases of sensory preconditioning procedures across species^1,4,11^. This region, which maintains continuous information exchange with other cortical areas, is thought to act as an integrator of past experiences into broader cognitive representations through a tightly regulated excitatory-inhibitory balance^15,16^. Previous studies have highlighted a specific role for hippocampal mechanisms during the encoding of incidental associations^11^. However, the contribution of specific hippocampal subregions or cell types to sensory preconditioning remains unclear. The dorsoventral axis of the rodent hippocampus is known to be structurally and functionally segregated. The dorsal hippocampus, connected to cortical regions and the thalamus, is primarily involved in cognitive processes such as navigation and exploration^17–21^. In contrast, the ventral hippocampus, which communicates with the amygdala, nucleus accumbens, and hypothalamus, plays a key role in motivated and emotional behaviors^17,18,22^. Based on this, our initial hypothesis was that the dorsal and/or ventral hippocampus may differentially contribute to sensory preconditioning in mice.

To test this, we employed chemogenetic and imaging techniques in combination with an adapted light-tone sensory preconditioning (_LT_SPC) procedure^6^. Our results suggest that the dorsal hippocampus, and in particular CaMKII-positive neurons, plays a crucial role in the formation of incidental associations between visual and auditory stimuli, ultimately driving sensory preconditioning responses. Understanding the differential contribution of the dorsal and ventral hippocampus to sensory preconditioning provides valuable insights into the mechanisms of higher-order conditioning and the broader role of the hippocampus in associative learning.

## Material and methods

### Animals

Male and female C57BL/6J mice were purchased from Charles River Laboratories and were used for behavioral and chemogenetic experiments. Male Parvalbumin (PV)-Cre mice were obtained by own breedings in our animal facility. All the mice used in this study were 8 weeks old at the beginning of the experiments and they were grouped-housed and maintained in a temperature (20–24°C) and humidity (40%–70%) controlled condition under a 12 h light/dark cycle and had ad libitum access to food and water. All behavioral studies were performed during the dark cycle (from 9am to 5pm) by trained researchers who were blind to the different experimental conditions.

All experimental procedures shown in this work have followed European guidelines (2010/62/EU) and were approved by the Committee on Animal Health and Care of Barcelona Biomedical Research Park (PRBB) and from the Generalitat de Catalunya. All experiments were performed in the animal facility of the PRBB, which has the full accreditation from the Association for Assessment and Accreditation of Laboratory Animal Care (AAALAC).

### Light-tone sensory preconditioning task (_LT_SPC)

The task was performed using automatized conditioning chambers (Imetronic, France) and was divided in different phases (Figure 1A):

**Figure 1.**
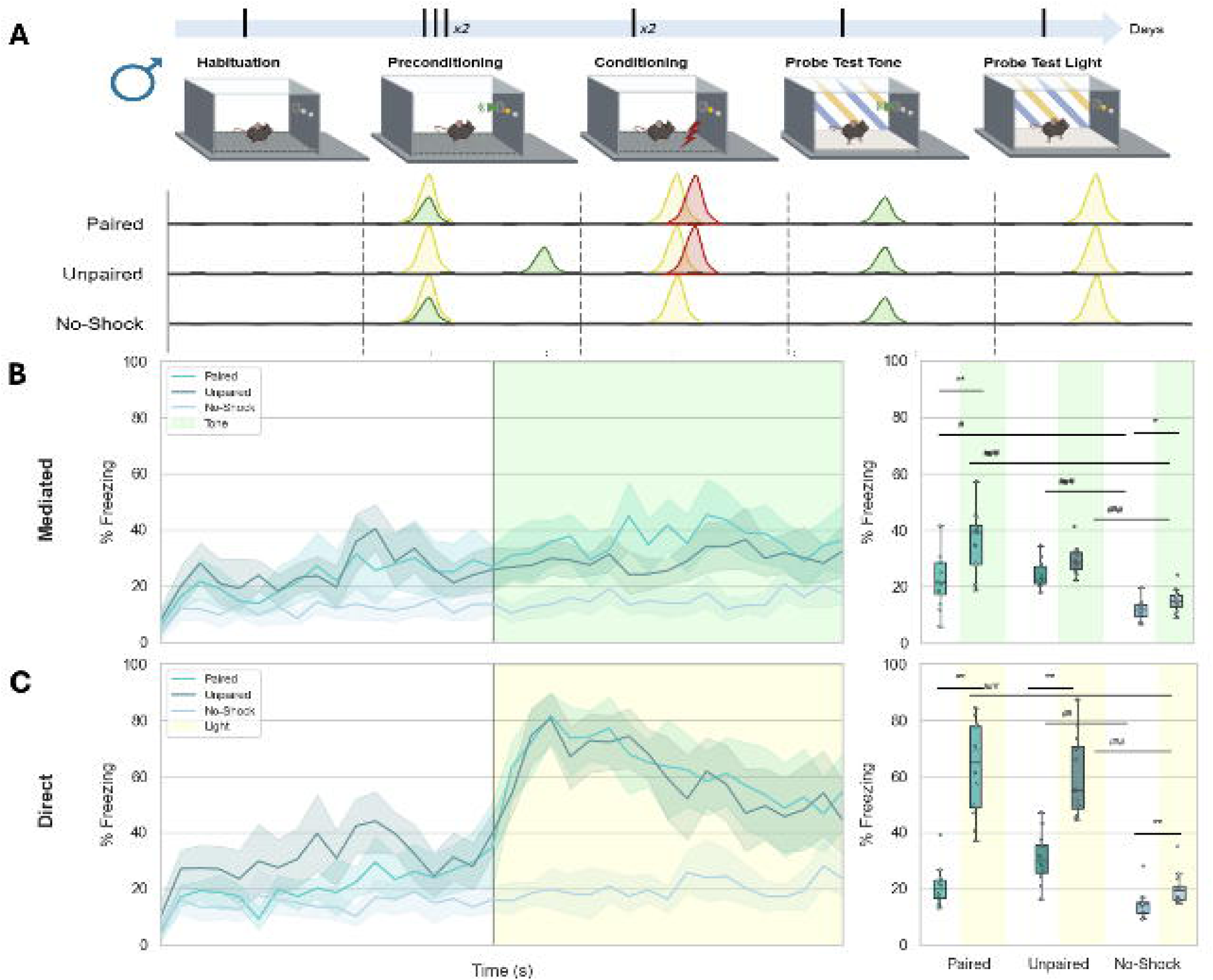
Simultaneous associations between light and tone are required for sensory preconditioning responding. (A) Schematic representation of the _LT_SPC task in males (2 training sessions) with a representation of paired, unpaired and no-shock experimental groups. The temporal dynamics of freezing represented in bins of 10 seconds across time of experiment of the Probe Test 1 (Tone) (B on the left) and Probe Test 2 (light) (C on the left). The percentage of time spent freezing during OFF and ON periods of the Probe Test 1 (Tone) (B on the right) and Probe Test 2 (light) (C on the right). * Significant p-value (<0.05) after false discovery rate (FDR). GLM: generalized linear model fitted to gamma distribution with planned comparisons. KW: Kruskal-Wallis across experimental groups: paired, unpaired, no-shock. See statistical details in Supplementary Table 1.

#### Habituation

Animals were exposed to the chambers with the presence of background noise (65db) (like white noise but allocated to different speakers to avoid that animals associate the sound with a location) during one session of 20 minutes.

#### Preconditioning Phase

This phase was composed by six sessions (2 per day; 3 hours of intersession interval) of 510 seconds each under background noise. These sessions started with an OFF period (only background noise) of 3 min and were followed by 5 simultaneous presentations of a light (CS1, white LED light of 1.8cm of diameter located in one side of the box) and a tone (CS2, 65db,3000Hz) during 30 seconds with an intertrial interval of 30 seconds and a final OFF period of 1 minute. This protocol details apply to the Paired and No-Shock experimental groups. For the unpaired groups, light and tone were never associated together during this phase. Indeed, animals were exposed to the same amount of time to light and tone but in different sessions (S1 light, S2 tone, S3 tone, S4 light, S5 light, S6 tone, in a pseudo-randomized way) separated by 3h (male and female mice) or 6h (female mice).

#### Conditioning Phase

This phase consisted of two training sessions in males and one training session in females. Each session lasted 510 seconds under background noise. In the case of males, there was an inter-session interval of 3 hours. Specifically, the sessions started with an off period of 3 min and were followed by 5 presentations of a 10-seconds light stimulus (CS1) that co-terminated during 2 seconds with a mild foot-shock (0.4 mA, see results) with an intertrial interval of 1 min and a final OFF period of 1 min. This protocol details apply to the Paired and Unpaired experimental groups. For the No-Shock group, animals followed the same conditioning session but without the exposure to the electric foot-shock. For the control group performed in female mice, no light was associated with the foot-shock during this conditioning session.

#### Probe Tests Phase

Mice were subjected on the same day to two Probe Test sessions where they were exposed to the tone (Mediated Learning, Probe Test 1) and the light (Direct Learning, Probe Test 2) in a new context (different from the previous phases). We changed 4 features of the context: floor texture, wall design, smell (from 70% ethanol to a CR36 disinfectant solution), and absence of background noise to avoid fear responses elicited by the context itself. The two Probe Tests lasted 6 minutes and were separated by at least 1 hour. In this session, animals underwent an OFF period of 3 min followed by an ON period of 3 min where the tone (Probe Test 1) or the light (Probe Test 2) were continuously exposed. All experimental groups (Paired, Unpaired and No-Shock) perform the Probe Tests in the same manner. The presence of mediated and direct learning is shown by the percentage of freezing during ON and OFF periods.

The behavior analysis was automatized using Deeplabcut to track the animal’s position across time and a Python homemade script to compute all the different analyses, graphs, and statistics performed in the present paper. The main behavior measured was the freezing response (i.e. immobility), which was defined as an Euclidean distance lower than 0.02 cm per pair of frames for videos with 25 fps, resulting in a speed lower than 0.5 cm/s. We validated this automated behavioral counting by performing correlations with manual counts and another software dedicated to freezing scoring (EzTrack)^23^, with which we found a high correlation (higher than 90%) (Supplementary Figure 1). The Deeplabcut project and all the scripts will be available in the GitHub repository of the lab.

### *In vivo* Fiber Photometry

Male C57BL/6J and PV-Cre mice were anesthetized with a mixture of ketamine (75 mg/Kg, Imalgene 500, Merial, Spain) and medetomidine (1 mg/Kg, Domtor, Spain) by intraperitoneal (i.p.) injection. Then, animals were placed into a stereotaxic apparatus (World Precision Instruments, FL, USA) with a mouse adaptor and lateral ear bars. For viral intra-hippocampus delivery, AAV vectors were injected with a Hamilton syringe coupled with a nanofill attached to a pump (UMP3-1, World Precision Instruments, FL, USA). PV-cre mice were injected with a mixture of 200 nl of AAV.Syn.NES.jRCaMP1a.WPRE.SV40 (titer: 1×10¹³, addgene 100848-AAV9) and 200nl of a Cre-dependent AAV-syn-FLEX-jGCaMP8f-WPRE (titer: 1×10¹³, addgene 162379-AAV9) in order to monitor both the activity of the synapsin-positive cells in red and the PV interneurons in green. C57BL/6J mice were injected with 400nl of AAV-CaMKII-GCaMP6m (titer: 1×10¹^2^, Tebu-BIO 189SL-AAV9) to monitor activity of the CaMKII positive neurons in green. To check DREADD functionality, 200 nl of the AAV-CaMKII-GCaMP6m and 200 nl of the inhibitory DREADD were infused in a similar manner. These viral vectors were infused (1 nl/s) directly into the dorsal (in one hemisphere) or ventral hippocampus (in the other hemisphere) with the following coordinates in mm: dorsal, AP ±1.5, ML ± 2, DV -1.5 and ventral, AP ±3.5, ML ± 3.3, DV -3.5, according to Paxinos and Franklin brain atlas (Paxinos and Franklin, 2001). After the AAV infusions, an optic Fiber (core 400 μm, N.A 0.5, RWD, China) was implanted using dental cement following the same coordinates except for DV that was placed 0.25 mm above viral infusions. Four weeks after this surgery, animals were used for *in vivo* calcium recordings where they undergo a _LT_SPC and *in vivo* recordings were performed during Light-Tone associations (preconditioning phase) and Light-Shock associations (conditioning phase). Before this experiment, animals were habituated for 3 days to connect and disconnect the optic fibers and to be habituated to the cable in the same chamber where the behavioral experiment was performed. During these two behavioral phases, *in vivo* recordings were performed using a commercial Fiber Photometry equipment (RWD, China) where we used 470nM LED to excite the GCaMP sensor, and 560 nM for the RCaMP signal. In all mice used, after the recording experiments, the signal was checked to validate the viral infusions.

To analyze the Fiber Photometry experiments, a custom python code was used and the behavioral videos and photometry recordings were synchronized by TTL signals. Raw calcium signals were pre-processed by removing the first minute of the recording to decrease the effect of the initial exponential photobleaching, and by removing point artifacts. The 470nM signal was fitted to the isosbestic 405nM using a linear fit and for each time point, ΔF/F was calculated as (F470nm –F405nm(fitted))/F405nm(fitted). This procedure is the same as described in previous works^24^. The codes used for the analysis will be found on the dedicated github repository.

### Chemogenetic modulation

Stereotaxic surgeries were performed as described above for fiber photometry. In this case, C57BL/6J or PV-cre mice were injected with 500 nl AAV-CaMKII-mCherry (titer: 7×10¹², 114469-AAV5) and 500 nl AAV-CaMKII-hM4Di (titer:7×10¹², 50477-AAV2) directly into the hippocampus, with the following coordinates: dorsal, AP ±1.5, ML ± 2, DV -1.5 and ventral, AP ±3.5, ML ± 3.3, DV -3.5, according to Paxinos and Franklin brain atlas (Paxinos and Franklin, 2001). On the other hand, PV-Cre mice were infused with AAV-DIO-hM4Di using the same coordinates to target the dorsal hippocampus. Three control groups were performed for each subregion (dorsal and ventral hippocampus): animals infused with control virus (pAAV-CaMKIIa-mCherry) and injected with saline or JHU37160 dihydrochloride (J60, 0.1 mg/Kg, i.p., HelloBio, HB6261)^25^, and animals infused with AAV-CaMKII-mCherry or AAV-DIO-hM4Di injected with saline. Animals were used for chemogenetics experiments four weeks after injections to get an optimal expression of the viruses. The injection of J60 was one hour before each preconditioning session (C57BL/6J and PV-Cre mice), before the Probe Test 1 (C57BL/6J mice) or the two conditioning sessions. In all mice used in the behavioral experiments, the signal was checked, and representative images are shown in Supplementary Figures.

### Histology

After the chemogenetic and imaging experiments, mice were anesthetized i.p with a mixture of ketamine (50 mg/Kg) and xylazine (20 mg/Kg) in overdose (3-4x body weight), transcardially perfused with cold 4% paraformaldehyde to fix tissues. Brains were removed and sectioned in serial coronal sections of 20 um and collected directly to the slide for further analysis in the microscope. All sections were counterstained with 4’,6-diamidino-2-phenylindole (DAPI, ref. 00-4959-52, Fluoromount-G w/DAPI, Life Technologies, USA) to visualize cell nuclei in the mounting medium. Slides were coverslipped and imaged by an epifluorescence Nikon Eclipse Ni-E. All animals were signal checked to guarantee the target of the hippocampus subregion.

### Data collection and statistical analysis

#### Data collection

All mice were randomly assigned to experimental conditions. Researchers performing the experiments were always blind to the condition of the subject and we used an automated way to analyze behavioral responses throughout the study. Raw data was processed and analyzed using homemade python packages.

#### Statistical analysis

Graphs and statistical analysis were performed through homemade python scripts that will be openly shared. All data comes from distinct mice and is shown as independent data points per animal ± standard error of the median (SEM). Normality and homoscedasticity of the data were assessed with the Kolmogorov-Smirnoff and Levene tests, respectively. Due to non-parametric properties of the data, a generalized linear model with gamma distribution was used for multivariable analysis (after a goodness of fit to select the appropriate distribution), followed by planned comparisons corrected with false discovery rate. For univariate analysis, the Kruskal-Wallis test was performed followed by planned comparisons corrected with false discovery rate. For simple comparisons the Mann-Whitney test was performed. Detailed statistical data for each experiment can be found in Extended Table 1 (for Main Figures) and Extended Table 2 (for Supplementary Figures).

## Results

### Simultaneous light-tone associations during preconditioning are required for sensory preconditioning responding in male mice

Sensory preconditioning tasks using olfactory and gustatory stimuli have already been demonstrated in mice in several previous studies^11,14^. However, other sensory modalities such as visual and auditory stimuli are also highly relevant for animals’ daily decision-making. While the use of these cues to assess aversive sensory preconditioning has been shown in rats^6,8,13^, their use in mice—particularly for aversive paradigms—remains limited^11^. The first aim of the present work was to establish a sensory preconditioning task in mice using auditory and visual cues, enabling the assessment of both mediated and direct learning. To this end, we developed a sensory preconditioning protocol (see Methods, Figure 1A), alongside a novel automated tool to analyze the freezing response (i.e., immobility) in mice. This tool combines DeepLabCut pose estimation with a newly developed Python script and its accuracy was validated showing a high correlation with manual scoring and ezTrack^23^ freezing assessments (Supplementary Figure 1). Although for freezing scoring our tool is similar to ezTrack measurements, there are several key added benefits offered by this approach such as an enhanced flexibility to combine freezing measurements with other customized behavioral assessments, which can be particularly beneficial for labs conducting large-scale or longitudinal behavioral studies in rodents.

Using both male and female mice, we conducted a long-term sensory preconditioning (_LT_SPC) experiment with three experimental groups in male (Figure 1A) and female mice (Supplementary Figure 2A): Paired group (protocol described in methods section); Unpaired group (same protocol, but the light and tone were never simultaneous); No-Shock group (same as the Paired group but without electric foot-shock exposure) (Figure 1A and Supplementary Figure 2A). According to this design, male (Figures 1B and 1C) and female mice (Supplementary Figure 2B and 2C) in the Paired group exhibited an enhanced freezing response at the onset of the tone or light, respectively. Specifically, exposure to the tone (Probe Test 1, ON period) led to increased freezing behavior (Figure 1B) compared to the OFF period, indicating sensory preconditioning responding in male mice. When the light was presented (Probe Test 2), animals showed greater freezing during the ON period compared to the OFF period (Figure 1C), revealing direct learning.

When the light and tone were temporally separated (Unpaired group), male mice failed to exhibit sensory preconditioning (Figure 1B), while their direct learning response to the light remained intact (Figure 1C). In contrast, female mice still displayed a behavioral response to the tone despite the separation of stimuli, which could be attributed to fear generalization (Supplementary Figure 2B), along with preserved direct learning (Supplementary Figure 2C). These results reveal sex differences in sensory preconditioning performance: male mice clearly demonstrated both mediated and direct learning, whereas female mice did not. To further explore this, we conducted an independent experiment to test whether the fear generalization observed in the female Unpaired group could be reduced by either extending the interval between cue presentations during preconditioning or by removing the light during the conditioning phase (Supplementary Figure 2D). Results showed that under both conditions, female mice still exhibited cue-induced fear generalization, which appears to mask their capacity for mediated learning responding.

Finally, in the No-Shock group, neither male (Figure 1B) nor female (Supplementary Figure 2B) mice showed a stimulus-mediated response, suggesting that tone or light exposure alone during the Probe Tests does not elicit behavioral changes. Altogether, these results confirm the successful establishment of an _LT_SPC protocol in male mice, which can now be used to further investigate the brain circuits involved in higher-order conditioning. However, additional research will be required to optimize an _LT_SPC protocol for female mice that avoids the confounding effect of fear generalization. Indeed, supporting these sex differences in this sensory preconditioning task, in male mice we observed a clear correlation between direct learning and mediated learning responding (Supplementary Figure 3A), whereas this is not shown in female mice (Supplementary Figure 3B).

### Hippocampal cells are engaged during _LT_SPC

Previous studies using sensory preconditioning paradigms with other sensory modalities have suggested a key role for the hippocampus in processing incidental associations between innocuous stimuli during the preconditioning phase^1,11^. Specifically, hippocampal GABAergic neurons expressing the type-1 cannabinoid receptors, but not the activity of PV interneurons, have been implicated in odor–taste associations in a mouse odor-taste sensory preconditioning task^26^. However, whether similar cellular mechanisms are involved in sensory preconditioning using different sensory modalities remains unknown. To better characterize the involvement of the dorsal and ventral hippocampus, as well as their specific cell types, during our _LT_SPC task, we performed simultaneous in vivo fiber photometry recordings by implanting an optic fiber in each hippocampal subregion of PV-Cre mice and infusing a mixture of AAV-Syn-RCaMP and AAV-DIO-GCaMP viruses (see *Methods*) in both hippocampal subregions. This allowed us to monitor general neuronal activity (via RCaMP) and the activity of PV-positive interneurons (via GCaMP) during both the preconditioning (light-tone pairings) and the conditioning phase (light-foot-shock pairings). Four weeks after surgery, animals were subjected to the _LT_SPC task (Figure 2A). Neuronal activity in the dorsal and ventral hippocampus was quantified as ΔF/F following stimulus pairings (dHPC: Figure 2B on left, vHPC: Figure 2C on left). Peri-event time histograms (PETHs) of Z-scored ΔF/F revealed the average dynamic response in each subregion during light–tone associations (dHPC: Figure 2B on right, vHPC: Figure 2C on right). These PETHs highlight a lower trial-by-trial variability in dHPC (Figure 2B) and a more sustained response over time compared to vHPC (Figure 2C). To quantify this activity, we measured the peak Z-scored ΔF/F within a 1-second window following stimulus onset and compared it to baseline values (Figure 2D). Notably, this increase in activity was consistent across all preconditioning sessions, with no significant differences observed when data were analyzed per session (Supplementary Figure 4A). Additionally, both neuronal activity markers (RCaMP and GCaMP) showed clear increases during the conditioning phase, both at the onset of the light stimulus (conditioned stimulus; Supplementary Figure 5) and at the time of foot-shock delivery (unconditioned stimulus; Supplementary Figure 6). These increases were consistent across sessions (Supplementary Figure 4B and C), with no significant session-dependent variation. Together, these findings demonstrate the engagement of hippocampal neurons, including PV-positive interneurons, in both the dHPC and vHPC during associative learning in our _LT_SPC task. This supports the role of the hippocampus in processing sensory associations involving non-olfactory modalities.

**Figure 2.**
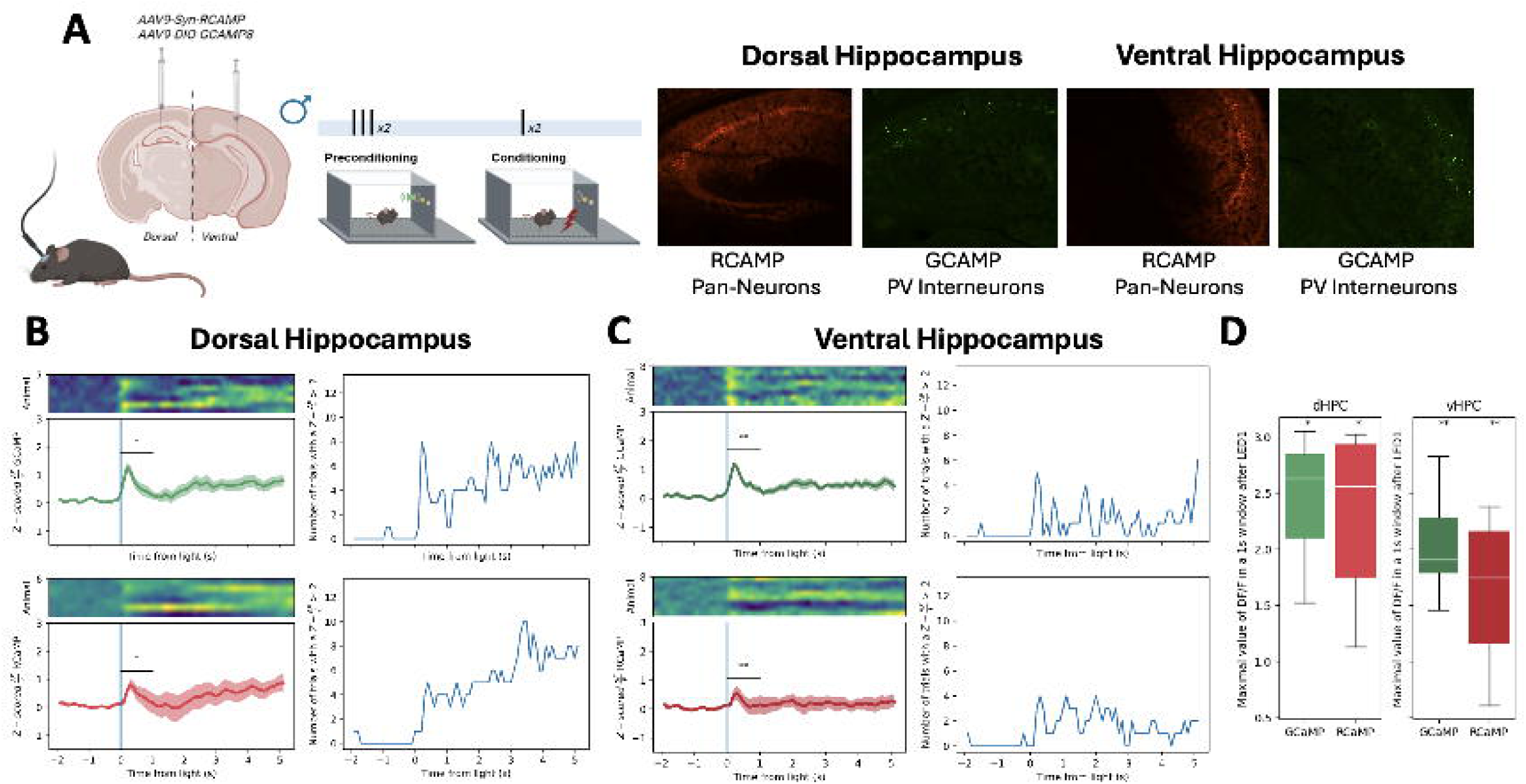
Hippocampal cell activity during _LT_SPC. A) Schematic representation of _LT_SPC task while fiber photometry recordings of RCaMP (calcium sensor in synapsin-positive neurons) and Cre-dependent GCaMP (calcium sensor in PV-positive interneurons) in dHPC and vHPC of PV^cre^ mice. B) dHPC modulation during preconditioning of _LT_SPC: on the left upper panel z scores of 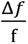 (where f represents fluorescence) of GCaMP PV-positive interneurons on dHPC (green), left bottom z scores of 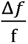 of RCaMP in neurons of dHPC (red), right upper number of events with a 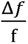 > 2 of GCaMP PV-positive interneurons on dHPC, and right bottom number of trials with a 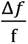 > 2 of Rcamp in neurons of dHPC. C) vHPC modulation during _LT_SPC: on left upper painel z scores of 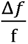 of GCaMP PV-positive inter neurons of vHPC (green), left bottom z scores of 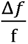 of RCaMP in neurons of vHPC (red), right upper number of trials with a 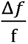 > 2 of Gcamp in PV-positive interneurons of vHPC, and right bottom number of trials with a 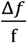 > 2 of RCaMP in neurons of vHPC. D) maximal value of 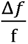in the first 1 second window after pairings compared with baseline, on the left (dHPC) and on the right (vHPC). * Significant p-value (<0.05). See statistical details in Supplementary Table 1.

Although our data showed a clear engagement of hippocampal activity at different phases of our _LT_SPC task, the use of a synapsin promoter is too broad to selectively point to a specific cell-type mediating the behavioral responses observed. Thus, we decided to use a similar approach and specifically monitor CaMKII+ neurons during the preconditioning sessions by infusing an AAV-CaMKII -GCAMP6 (see Methods, Figure 3A). Interestingly, and matching what was observed with a pan promoter (*i.e.* synapsin), PETHs of Z-scored ΔF/F (Figure 3B and C) and the subsequent histograms representing the peak Z-scored ΔF/F within a 1-second window following stimulus onset (Figure 3D) revealed a clear engagement of CaMKII-positive neurons from both hippocampal subregions during light–tone associations, which is constant across preconditioning sessions (Figure 3E).

**Figure 3.**
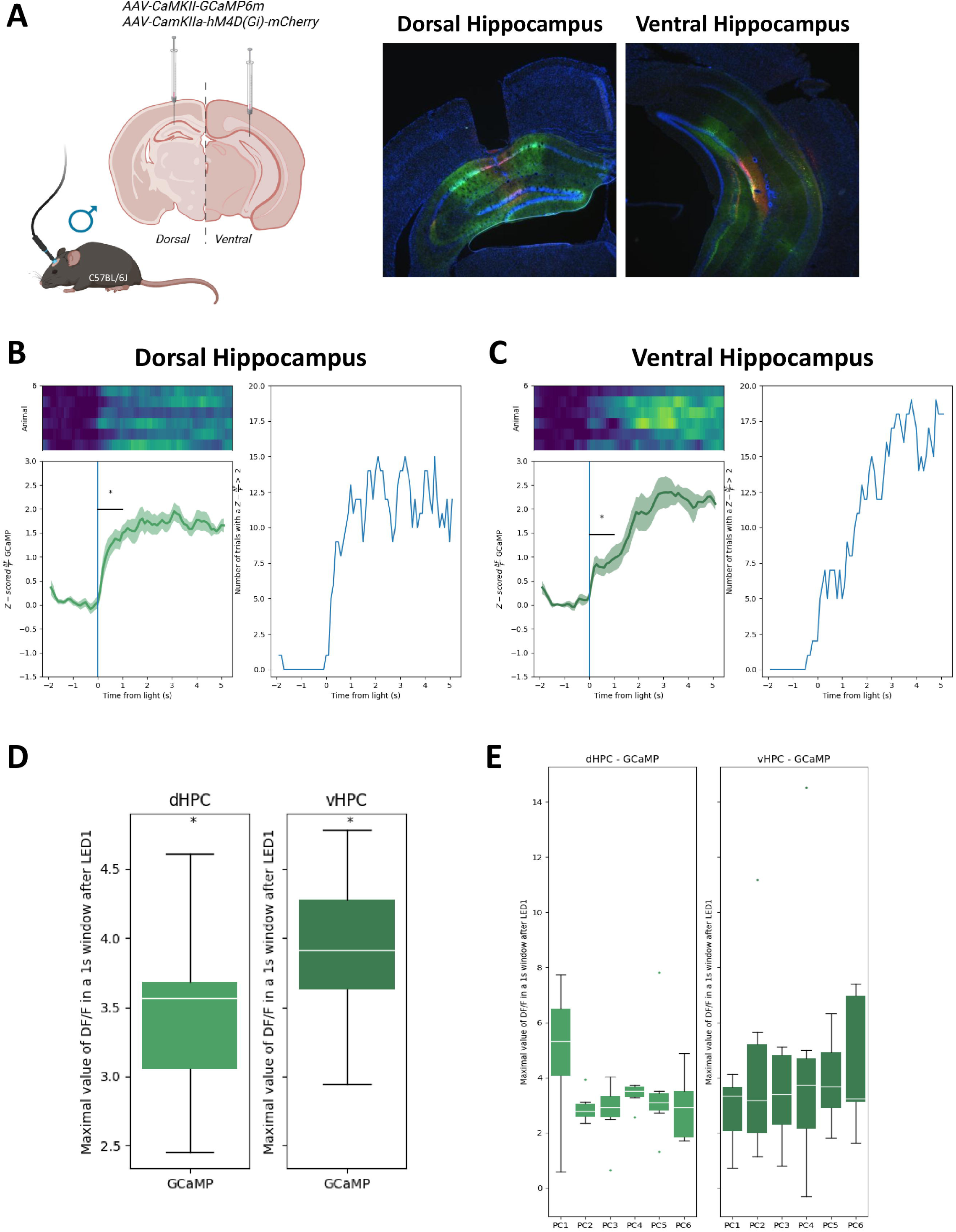
CaMKII neurons during preconditioning on dHPC and vHPC. A) maximal value of 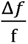 in the first 1 second window after stimulus compared with baseline, for each stimulus during preconditioning session, on the left (dHPC) and on the light (vHPC). B) dHPC modulation during preconditioning of _LT_SPC: on left z scores of 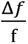 (where f represents fluorescence) of GCaMP in CaMKII-positive neurons on dHPC (green), on right number of events with a 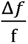 > 2 of GCaMP CaMKII-positive neurons on dHPC. C) vHPC modulation during preconditioning of _LT_SPC: on left z scores of 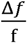 (where f represents fluorescence) of GCaMP in CaMKII-positive neurons on vHPC (green), on right number of events with a 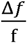 > 2 of GCaMP CaMKII-positive neurons on vHPC. D) maximal value of 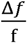 in the first 1 second window after stimulus compared with baseline, for the average of the 6 pairings stimuli during preconditioning session, on the left (dHPC) and on the light (vHPC). *, p-value (<0.05). Statistical details in Supplementary Table 2.

### CaMKII-positive neurons in dorsal hippocampus mediate _LT_SPC

The in vivo photometry results suggest a prominent engagement of the dorsal hippocampal (dHPC) subregion, characterized by increased activity of various hippocampal cell types, including CaMKII-positive neurons, during light–tone presentations in the preconditioning phase. However, no study to date has explored a potential causal dissociation between the roles of the dHPC and vHPC in mouse sensory preconditioning. To investigate this, adeno-associated viral vectors expressing an inhibitory DREADD (hM4DGi, hereafter referred to as DREADD-Gi)^27^ under the CaMKII promoter were infused into either the dHPC or vHPC to selectively inhibit principal excitatory neurons in these hippocampal subregions (Figure 4A). Control animals, which received either saline or the DREADD agonist J60, were pooled as they exhibited no significant behavioral differences and showed a reliable mediated and direct learning in the _LT_SPC task (Figure 4B and, Supplementary Figure 7A-D).

**Figure 4.**
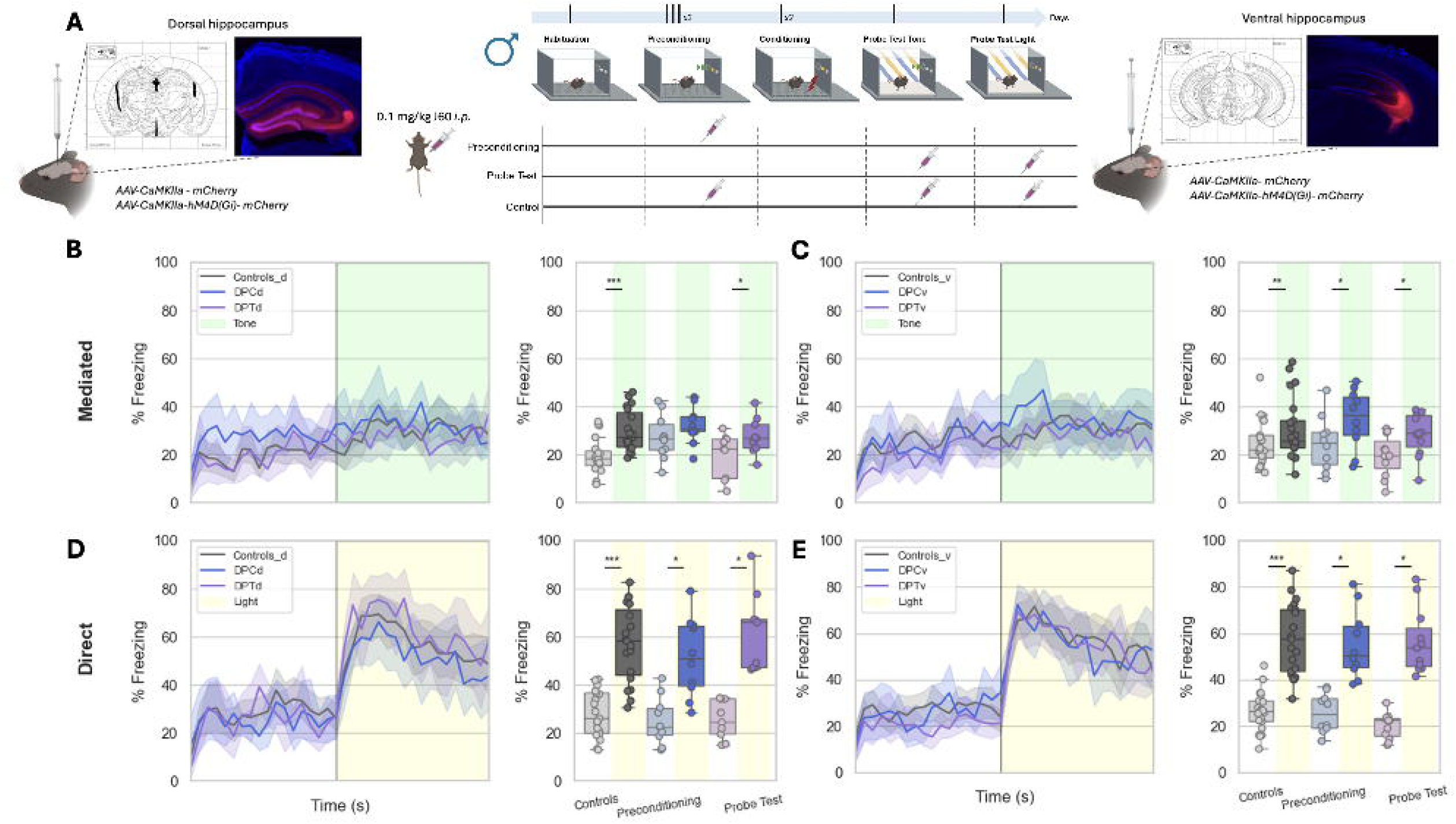
Chemogenetic modulation of dorsal and ventral hippocampus during _LT_SPC. (A) Schematic representation of _LT_SPC task combined with chemogenetic approaches, where an inhibitory DREADD was infused in dHPC (representative image on the left) and vHPC (representative image on the right). During the preconditioning (DPC) or the Probe Test (DPT), a DREADD agonist (J60) was injected intraperitoneally (controls where injected with the agonist in both phases) (diagram of DREADD agonist administration on the center). (B,C,D,E on the left) The temporal dynamics of freezing represented in bins of 10 seconds across time of experiment of the Probe Test 1 (Tone) of dHPC (B on the left) and vHPC (C on the left) and Probe Test 2 (light) of dHPC (D on the left) and vHPC (E on the left). (B, C on the right) The percentage of time spent in freezing during OFF and ON periods (left) of the Probe Test 1 (Tone) of animals infused in dHPC (B on the right) and vHPC (C on the right) D, E). (D, E on the right) The percentage of time spent in freezing during OFF and ON periods (left) of the Probe Test 2 (Light) of animals infused in dHPC (D on the right) and vHPC (E on the right). Controls are labeleld as Constrols_d (on dHPC) and Controls_V (on vHPC). The preconditioning groups are labelled as DPCd (on dHPC) and DPCv (on vHPC). The Probe Test groups are labelled as DPTd (on dHPC) and DPTv (on vHPC). *Significant p-value (<0.05) after false discovery rate (FDR). GLM: generalized linear model fitted to gamma distribution with planned comparisons. See statistical details in Supplementary Table 1.

Notably, inhibition of CaMKII-positive neurons in the dHPC, which decrease calcium activity in CaMKII-positive neurons (Supplementary Figure 7E and F), during the preconditioning phase (i.e., J60 administration in DREADD-Gi mice) completely abolished mediated learning (Figure 4B) without affecting direct learning (Figure 4D). This effect was specific to the preconditioning phase, as inhibition prior to conditioning sessions (Supplementary Figure 7G) or before Probe Test 1 (Figure 4B) did not impair mediated learning. In contrast, inhibition of CaMKII-positive neurons in the vHPC, whether during preconditioning or Probe Test 1, had no effect on either mediated (Figure 4C) or direct learning (Figure 4E). These findings indicate that the dHPC, but not the vHPC, plays a specific and causal role in encoding associations between innocuous stimuli (e.g., light and tone) that underlie sensory preconditioning responses. Given the additional results observed in our photometry experiments and the crucial role of dHPC, we next investigated whether PV-positive interneurons, which were also activated during stimuli presentation in our photometry recordings, played a role in this behavior. To test this, a Cre-dependent DREADD-Gi virus was infused into the dHPC of PV-Cre mice to allow for selective inhibition of PV interneurons (Supplementary Figure 8A-C) during light–tone pairings. Remarkably, silencing PV interneurons during the preconditioning phase did not affect either mediated or direct learning (Supplementary Figure 8D).

Taken together, these results suggest that CaMKII-positive principal neurons, but not PV interneurons, in the dHPC, are critical for encoding light–tone associations that drive sensory preconditioning responses in mice. Importantly, no statistical behavioral differences were observed during the conditioning across the different experiments (Supplementary Figure 9).

## Discussion

Our study provides novel insights into the role of the hippocampus in higher-order conditioning by establishing and validating a light-tone sensory preconditioning (_LT_SPC) task in male mice. Using in vivo calcium imaging, we linked the activity of specific hippocampal cells in the dorsal and ventral subregions to distinct behavioral phases of the task. Furthermore, chemogenetic inhibition experiments allowed us to causally identify the dorsal hippocampus (dHPC)—but not the ventral hippocampus (vHPC)—as essential for encoding light–tone pairings, which are critical for the behavioral outcomes observed in sensory preconditioning.

Sex differences in classical fear conditioning have been reported, with conflicting findings showing stronger conditioning in males, in females, or no differences at all^28–35^. However, to our knowledge, no previous studies have examined sex differences in sensory preconditioning paradigms. Our results demonstrate that, using the same preconditioning and conditioning settings, only male mice show a reliable light-tone sensory preconditioning responding, whereas female mice show behavioral responses mostly associated to fear generalization. Indeed, our results could suggest that female mice are more vulnerable to aversive stimuli responding, which might be consistent with prior evidence showing sex-dependent differences in conditioned freezing responses^36–39^. When the light and the tone were presented separately during the preconditioning, or when the foot-shock was omitted, both mediated and direct learning were abolished in male mice. Instead, female mice still showed reduced but detectable cue-induced responses under these conditions, even if we increased the time between light and tone exposures during the preconditioning phase. Thus, we did not successfully set up a light-tone sensory preconditioning paradigm for female mice and future research will be required to optimize and adapt a protocol to reliably observe mediated learning in females. Given that female mice also tend to exhibit greater fear generalization^29,40,41^, the observed differences in the unpaired group could reflect this enhanced generalization^33^. In this regard, changes on the number of conditioning sessions and/or trials, intensity of the electric foot-shock or additional contextual changes could avoid this fear generalization in future sensory preconditioning paradigms in female mice. In addition, an interesting unresolved issue is how sex hormones could alter sensory preconditioning responding or the different performance of male and female mice in different protocols. Overall, our data clearly revealed the necessity to adapt behavioral protocols for each sex in sensory preconditioning paradigms involving auditory and visual stimuli and aversive conditioning.

Prior research has established the hippocampus as critical for encoding spatial location, context, cues, and memory related to appetitive and aversive behaviors^42–46^. However, few studies have addressed its causal involvement in reinforcement learning^47^, and even fewer have explored its role in forming associations between innocuous stimuli. Using in vivo fiber photometry, we observed increased neuronal activity in both the dorsal and ventral hippocampus during light–tone pairings and during light–shock associations. These findings support the notion that the hippocampus is actively engaged in encoding both conditioned (CS) and unconditioned (US) stimuli, in agreement with studies showing hippocampal activation during reinforcement behavior^47–49^. Still, some prior work has argued that the hippocampus is not required for forming simple cue–US associations, which may instead rely on the amygdala^50^.

Our photometry results show that the simultaneous presentation of sensory cues is encoded by the hippocampus, reinforcing its role in associative learning and sensory integration^51^. This is consistent with previous work in both animals and humans supporting hippocampal involvement during associative phases of sensory preconditioning^1,11,13,14,52^. Importantly, however, none of these earlier studies investigated the specific roles of hippocampal subregions or cell types. Our data indicate that modulation of hippocampal activity—specifically in the dHPC—affects encoding during the preconditioning phase but not during conditioning or retrieval phases. Chemogenetic inhibition of CaMKII-positive neurons in the dHPC during the preconditioning blocked mediated learning responding, while inhibition in the vHPC had no such effect. Moreover, silencing PV-positive interneurons in the dHPC during preconditioning did not disrupt sensory preconditioning, suggesting that principal excitatory neurons, but not PV interneurons, are essential for forming associations between innocuous stimuli. Indeed, it is somehow contradictory to observe an activity enhancement of PV-positive interneurons during light-tone associations and the lack of effect of its inhibition during this phase. To fully discard any role of the activity of these interneurons during this phase, future experiments should aim to particularly inhibit these cells during restricted temporal windows such as some preconditioning sessions rather than the full preconditioning phase or to specifically inhibit its activity during the light-tone exposures (e.g. close-loop optogenetic approaches).

Other brain regions, including the orbitofrontal, retrosplenial, and perirhinal cortices, have also been implicated in processing associations between neutral sensory cues^6–8,10^. These areas maintain reciprocal anatomical connections with the hippocampus^53–55^, suggesting a broader, hippocampus-guided network for encoding higher-order associations. Our findings showing that the dHPC—but not the vHPC—is crucial for encoding light–tone pairings may offer new insight into this circuit. A likely candidate for mediating such interactions is the retrosplenial cortex, which receives direct projections from the dHPC^56^. Thus, inhibiting CaMKII-positive neurons in the dHPC, which project to the retrosplenial cortex could disrupt sensory preconditioning, supporting this potential circuit model. Future studies will be necessary to explore this hypothesis in detail.

Despite the robust findings presented by our data, several limitations must be acknowledged. First, while we successfully validated the _LT_SPC protocol in male mice, further optimization is needed for its reliable use in female mice as commented above. Second, the fixed order of Probe Tests may have introduced extinction effects for the tone^57^, potentially affecting responses to the light cue. Future studies should counterbalance Probe Tests to address this possibility. Third, our behavioral protocol required context changes between experimental phases, which may have engaged hippocampal circuits^58^ more strongly than if a single context had been used. Thus, the impact of context-switching on hippocampal involvement should be addressed in future work. Finally, this work has been focused on the activity of CaMKII-positive neurons and PV-interneurons although we cannot discard the role of other hippocampal cell types such as somatostatin interneurons, which have been recently involved in learning and associative processes^59,60^. Future experiments will be required to fully characterize the role of different hippocampal subtypes in sensory preconditioning. Indeed, other techniques such as the optogenetic modulation of cell activity could give more details on the temporal engagement of these hippocampal circuits in higher-order conditioning.

In summary, our results identify a key role for hippocampal circuits—particularly CaMKII-positive neurons in the dorsal hippocampus—in mediating associations between innocuous stimuli during light-tone sensory preconditioning. This process reflects a fundamental cognitive mechanism underpinning higher-order learning and decision-making in both animals and humans. Understanding the neural substrates of sensory preconditioning is crucial, as this form of learning may contribute to adaptive behavior but also underlie maladaptive processes seen in disorders such as psychosis^61,62^.

## Supporting information

Supplementary Figures and statistical Tables

## Acknowledgements

We would like to thank the personnel of the Animal Facility of the Parc de Recerca Biomedica de Barcelona (PRBB) for mouse care. We thank all the members of our lab for useful discussions during the development of the project and Remi Proville (Aquineuro, Bordeaux, France) for the great help in the analysis of behavior and *in vivo* photometry. Finally, we would like to also thank Dr. Maria Victoria Puig Velasco for providing the PV-Cre colony. This work was supported by la Generalitat de Catalunya (SGR (00022) and “Jo Investigo” (2022 INV-1 00005/100005TG3) programmes) from the Departament d’Economia i Coneixement de la Generalitat de Catalunya (Spain) and from the European Research Council (ERC) under the European Union’s Horizon 2020 research and innovation programme (Grant agreement No. [948217]).

## Author Contributions

A.B-G. and J.P. contributed to the conception of the project and performed and analyzed all the experiments. C.R-D. helped on all surgical procedures and immunohistochemistry assays. I.M-S. helped with behavioral and immunohistochemistry assays. A.B-G. and J.P wrote the manuscript. All authors have corrected and revised the manuscript.

