## Supplementary Figures and statistical Tables for "Dorsal hippocampus mediates light-tone associations in male mice"

[2] Current address: Gulbenkian Institute for Molecular Medicine, Oeiras, Portugal

**Content:**

**Supplementary Figures 1-9**

**Supplementary Tables 1-2**

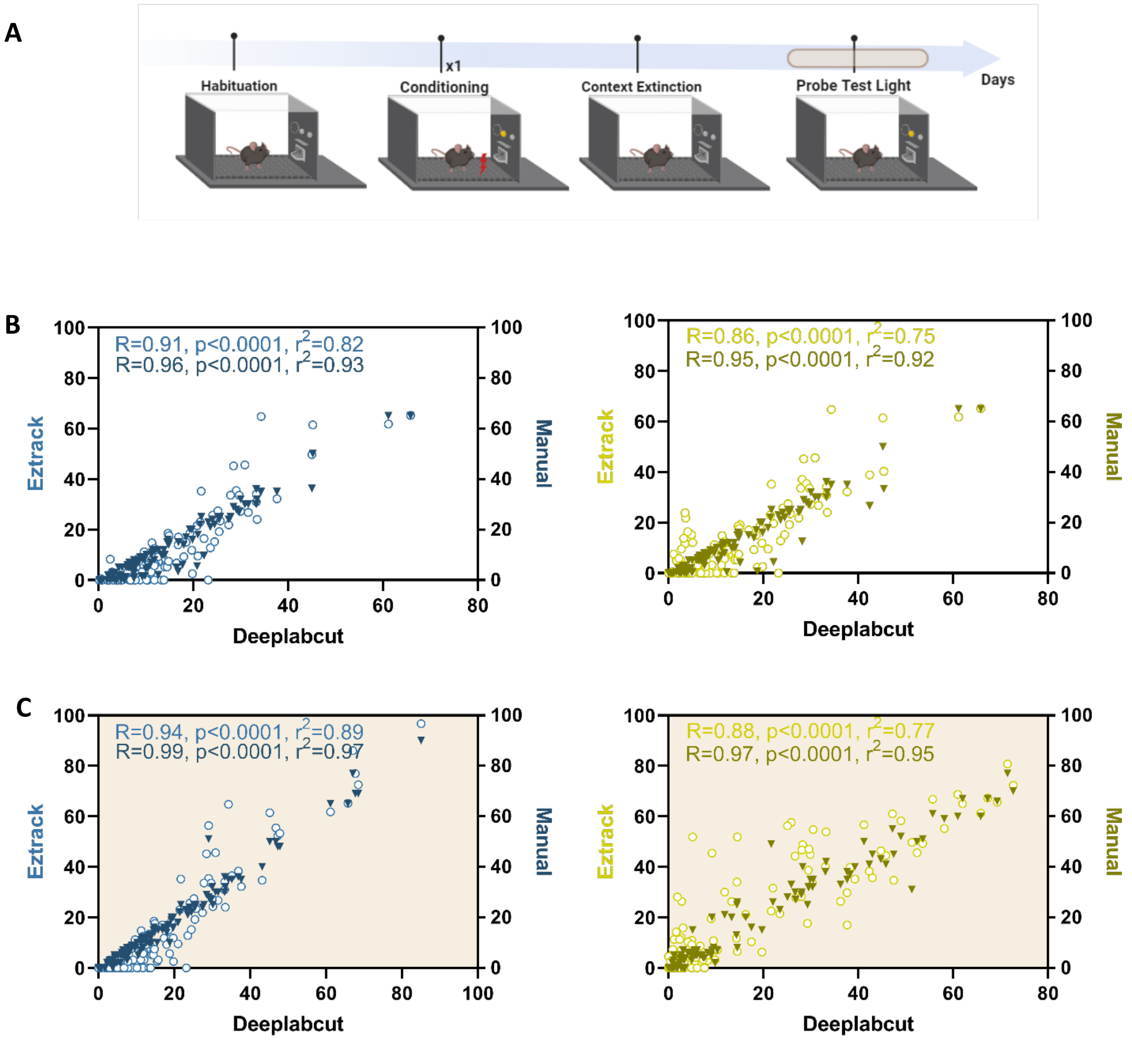


**Supplementary Figure 1. New automatized tool to measure freezing responses.** We developed a new automatized tool to assess Freezing behavior using a fear conditioning protocol (A). This novel tool was based on pose estimation using DeepLabCut and a customized python script, which considers freezing when the euclidean distance is lower than 0,02 cm per pair of frames for videos with 25 fps, resulting in a speed lower than 0,5 cm/s. To validate this tool, we correlate the freezing measures during all phases (upper) (B) and probe test (lower) (C) on males (left) and on females (right) with ezTrack (a software dedicated to the automatized freezing quantification) and our own manual counting performed by an expert in this type of behaviors. See Statistical details in Supplementary Table 2.

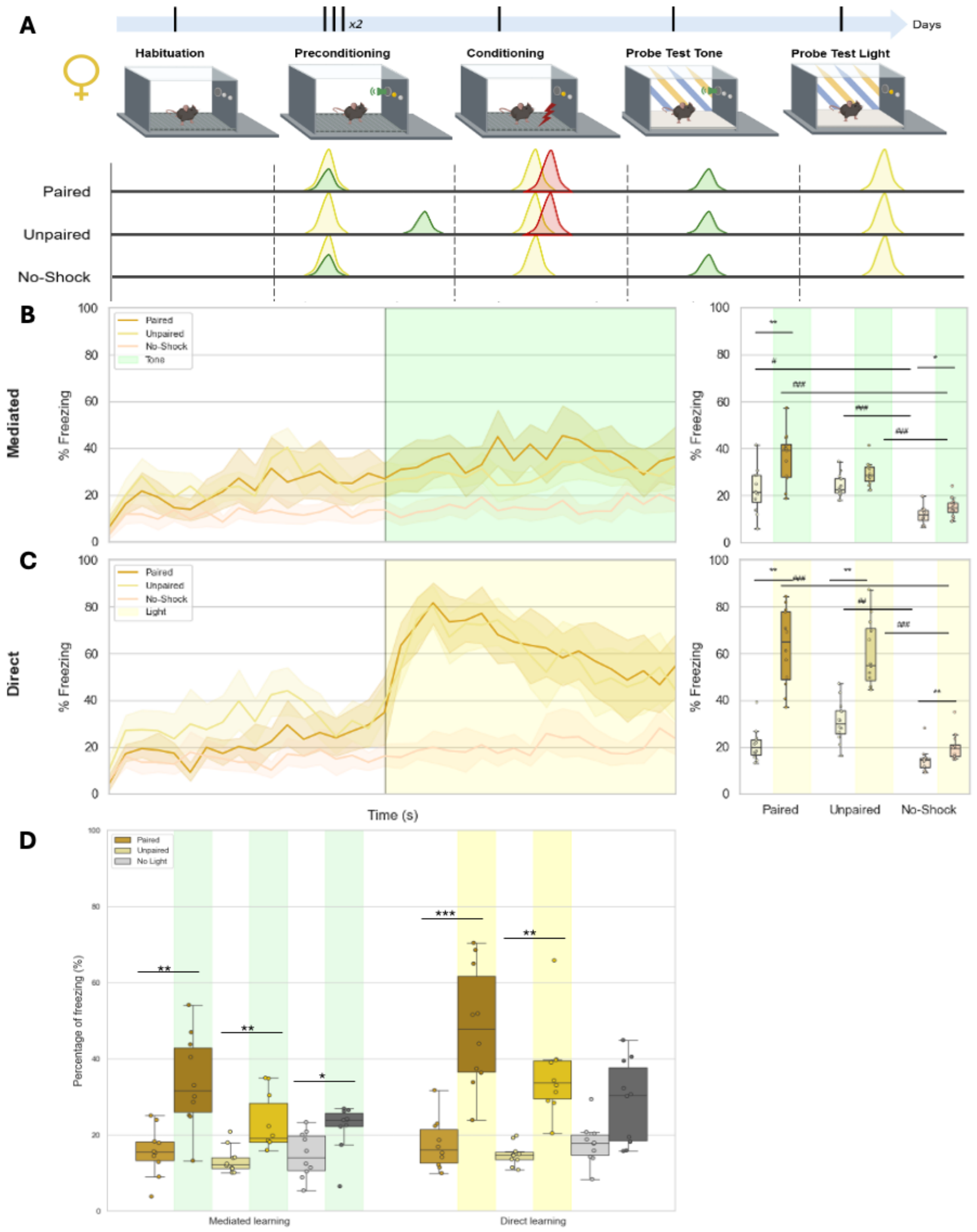


**Supplementary Figure 2. Simultaneous associations between light and tone to develop a sensory preconditioning protocol in females.** (A) Schematic representation of the $_{LT}$SPC task in females (1 training sessions) with a representation of paired, unpaired and no-shock experimental groups. The temporal dynamics of freezing represented in bins of 10 seconds across time of experiment of the probe test 1 (Tone) (B on the left) and probe test 2 (light) (C on the left). The percentage of time spent freezing during OFF and ON periods of the probe test 1 (Tone) (B on the right) and probe test 2 (light) (C on the right). (D) Independent experiment to test fear generalization observed in the female, we extended the interval between cue presentations during preconditioning (Unpaired) or by removing the light during the conditioning phase (No light).* Significant p-value (<0.05) after false discovery rate (FDR). GLM: generalized linear model fitted to gamma distribution with planned comparisons. See statistical details in Supplementary Table 1.

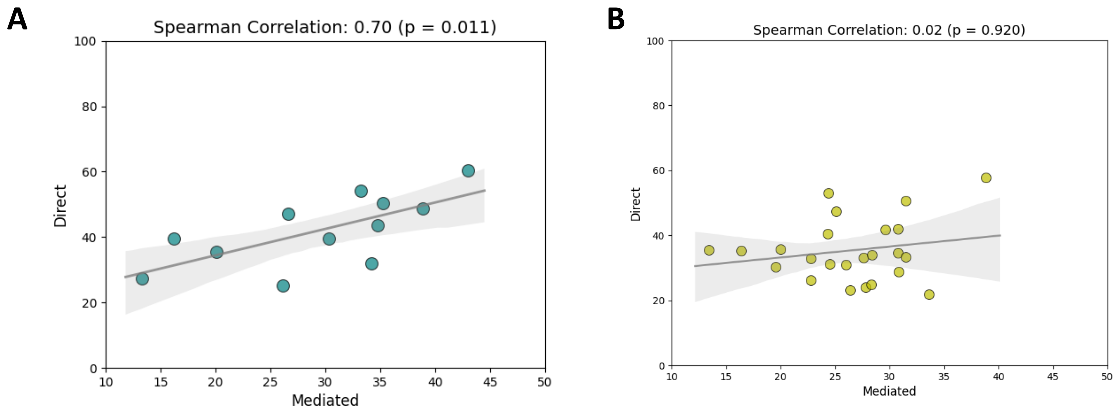


**Supplementary Figure 3. Spearman Correlations of mediated (Tone) and direct (Light) learning for the protocol of sensory preconditioning for males (A) and females (B).** Panel A shows a significant positive correlation in males (ρ = 0.70, $p$ = 0.011), whereas panel B shows no significant correlation in females (ρ = 0.02, $p$ = 0.920).

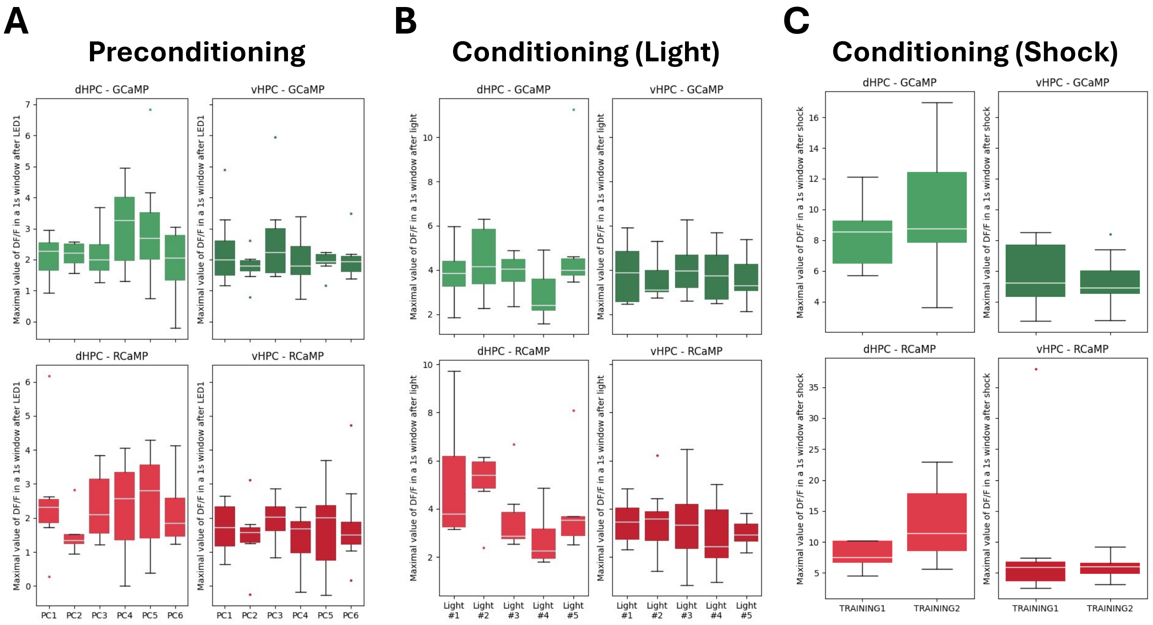


**Supplementary Figure 4.** Fiber photometry recordings of RCaMP (calcium sensor in synapsin-positive neurons) and Cre-dependent GCaMP (calcium sensor in PV-positive interneurons) in dHPC and vHPC of $PV^{cre}$ mice during each preconditioning trial (A), each light presentation during conditioning (B) and each shock presentation during conditioning (C). A,B,C maximal value of $\frac{\Delta f}{f}$ in the first 1 second window after stimulus compared with baseline, on the upper (GCaMP) and on the lower (RCaMP) in dHPC and vHPC. * Significant p-value (<0.05). See statistical details in Supplementary Table 1.

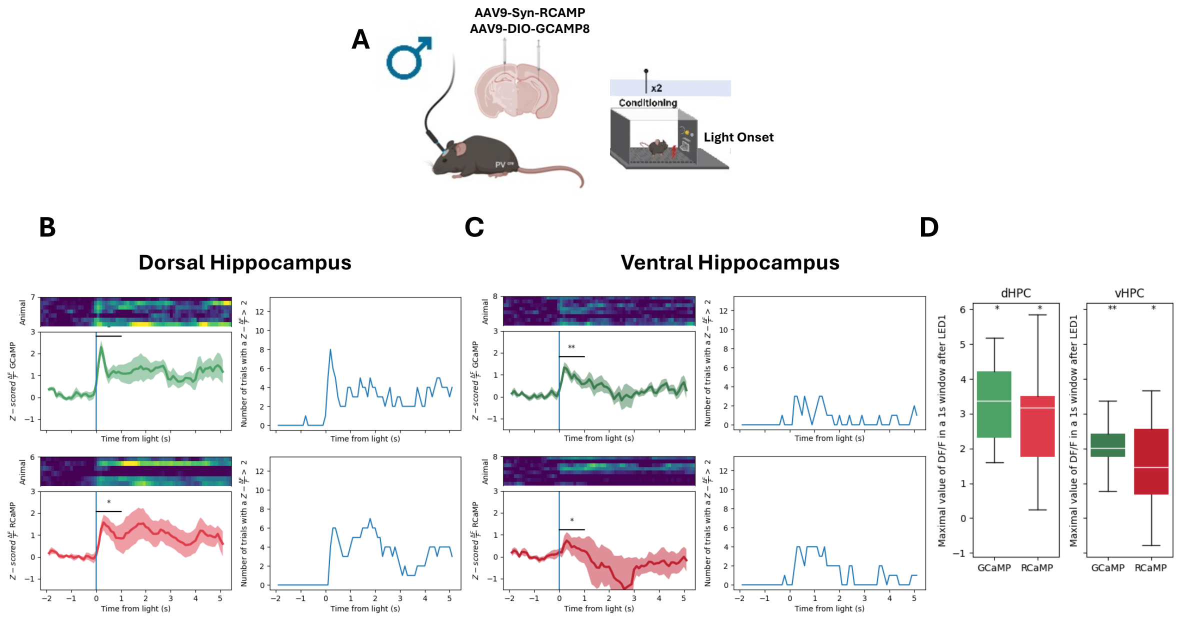


**Supplementary Figure 5. Hippocampal cells activity at the onset the light in conditioning sessions.** On the top, schematic representation of fiber photometry recordings with RCAMP1 and GCAMP8 in dHPV and vHPC of PV-Cre mice at the onset of the conditioned light (i.e. after pairing between light and electric footshock) during the conditioning phase. B) dHPC modulation during preconditioning of $_{LT}$SPC: on left upper panel, z scores of $\frac{\Delta f}{f}$ (where f represents fluorescence) of GCAMP8 PV-positive interneurons on dHPC (green); left bottom panel, z scores of $\frac{\Delta f}{f}$ of RCAMP1 neurons in dHPC (red); right upper panel, number of events with a $\frac{\Delta f}{f} > 2$ of GCAMP8 PV-positive interneurons in dHPC, and right bottom panel, number of trials with a $\frac{\Delta f}{f} > 2$ of RCAMP1 neurons in dHPC. C) vHPC modulation during $_{LT}$SPC: on left upper panel, z scores of $\frac{\Delta f}{f}$ of GCAMP8 PV-positive interneurons in vHPC (green); left bottom panel, z scores of $\frac{\Delta f}{f}$ of RCAMP1 neurons in vHPC (red); right upper panel, number of trials with a $\frac{\Delta f}{f} > 2$ of GCAMP8 PV-positive interneurons in vHPC, and right bottom panel, number of trials with a $\frac{\Delta f}{f} > 2$ of RCAMP1 neurons in vHPC. D) maximal value of $\frac{\Delta f}{f}$ in the first second window after pairings compared with baseline, on left (dorsal hippocampus, dHPC) and on right (ventral hippocampus, vHPC). *, p-value (<0.05). Statistical details in Supplementary Table 2.

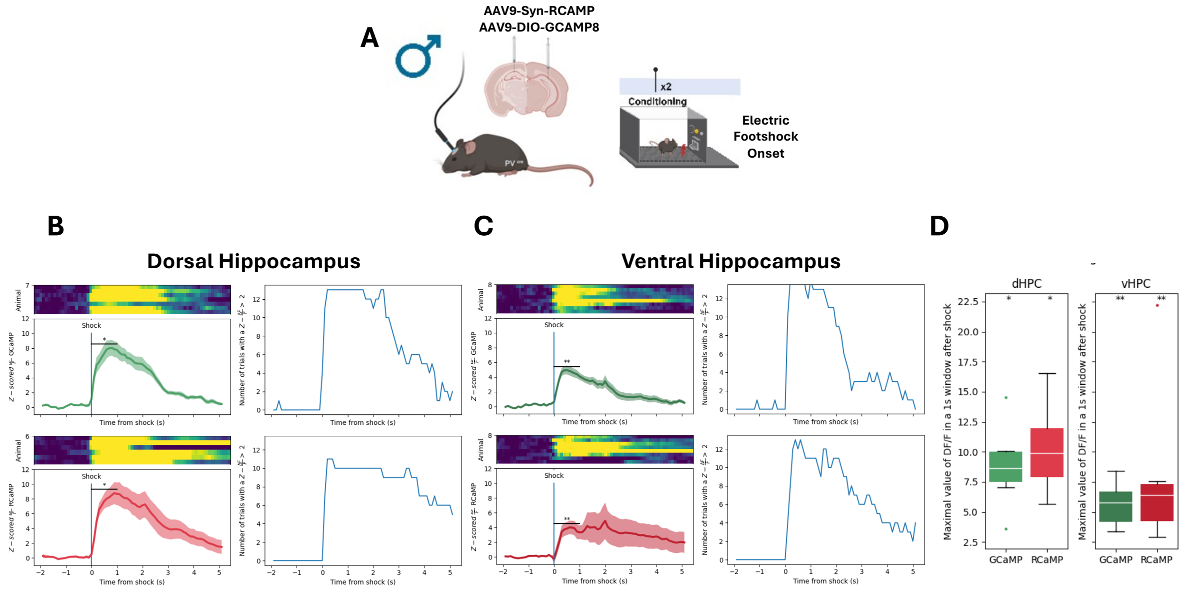


**Supplementary Figure 6. Hippocampal cells activity at the electric footshock onset.** On the top, schematic representation of the fiber photometry recordings with RCAMP1 and GCAMP8 on dorsal or ventral hippocampus of PV-Cre mice at the onset of the electric footshock during the conditioning phase. B) B) dHPC modulation during preconditioning of ${}_{LT}$SPC: on left upper panel, z scores of $\frac{\Delta f}{\mathrm{f}}$ (where f represents fluorescence) of GCAMP8 PV-positive interneurons on dHPC (green); left bottom panel, z scores of $\frac{\Delta f}{\mathrm{f}}$ of RCAMP1 neurons in dHPC (red); right upper panel, number of events with a $\frac{\Delta f}{\mathrm{f}} > 2$ of GCAMP8 PV-positive interneurons in dHPC, and right bottom panel, number of trials with a $\frac{\Delta f}{\mathrm{f}} > 2$ of RCAMP1 neurons in dHPC. C) vHPC modulation during ${}_{LT}$SPC: on left upper panel, z scores of $\frac{\Delta f}{\mathrm{f}}$ of GCAMP8 PV-positive interneurons in vHPC (green); left bottom panel, z scores of $\frac{\Delta f}{\mathrm{f}}$ of RCAMP1 neurons in vHPC (red); right upper panel, number of trials with a $\frac{\Delta f}{\mathrm{f}} > 2$ of GCAMP8 PV-positive interneurons in vHPC, and right bottom panel, number of trials with a $\frac{\Delta f}{\mathrm{f}} > 2$ of RCAMP1 neurons in vHPC. D) maximal value of $\frac{\Delta f}{\mathrm{f}}$ in the first second window after pairings compared with baseline, on left (dorsal hippocampus, dHPC) and on right (ventral hippocampus, vHPC). *, p-value (<0.05). Statistical details in Supplementary Table 2.

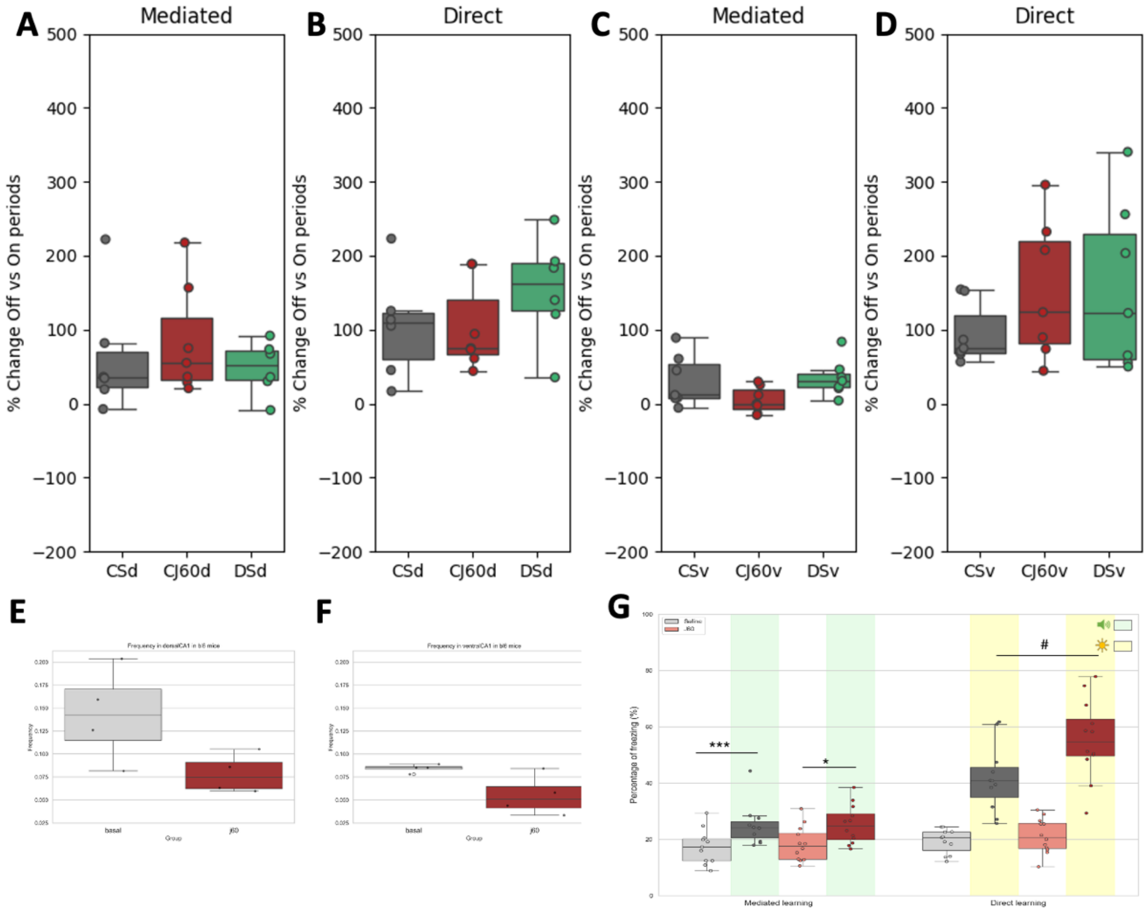


**Supplementary Figure 7. Additional controls for chemogenetics experiments.** A,B,C,D percentage change of the Off versus On periods to the control groups during mediated (A) and direct (B) of dHPC and mediated (C) and direct (D) of vHPC. The controls groups are constituted by: CS (animals injected in HPC with AAV-CamkII-mCherry with saline (i.p.) during preconditioning and probe test), CJ60 (animals injected in HPC with AAV-CamkII-mCherry with J60 (i.p.) during Preconditioning and probe test), and DS (animals injected in HPC with AAV-CamkII-hM4Di with J60 (i.p.) during Preconditioning and probe test). E) Frequency of calcium transients in CaMKII-positive neurons in the dHPC in C57BL6J mice in basal and J60 conditions. F) Frequency of calcium transients in CaMKII-positive neurons in the vHPC in C57BL6J mice in basal and J60 conditions. G) During the conditioning phase, the DREADD agonist J60 was injected intraperitoneally and the percentage of time spent in freezing during OFF and ON periods of the Probe Test 1 (Mediated learning) and Probe Test 2 (Direct learning) is shown. **, p-value (<0.01). Statistical details in Supplementary Table 2.

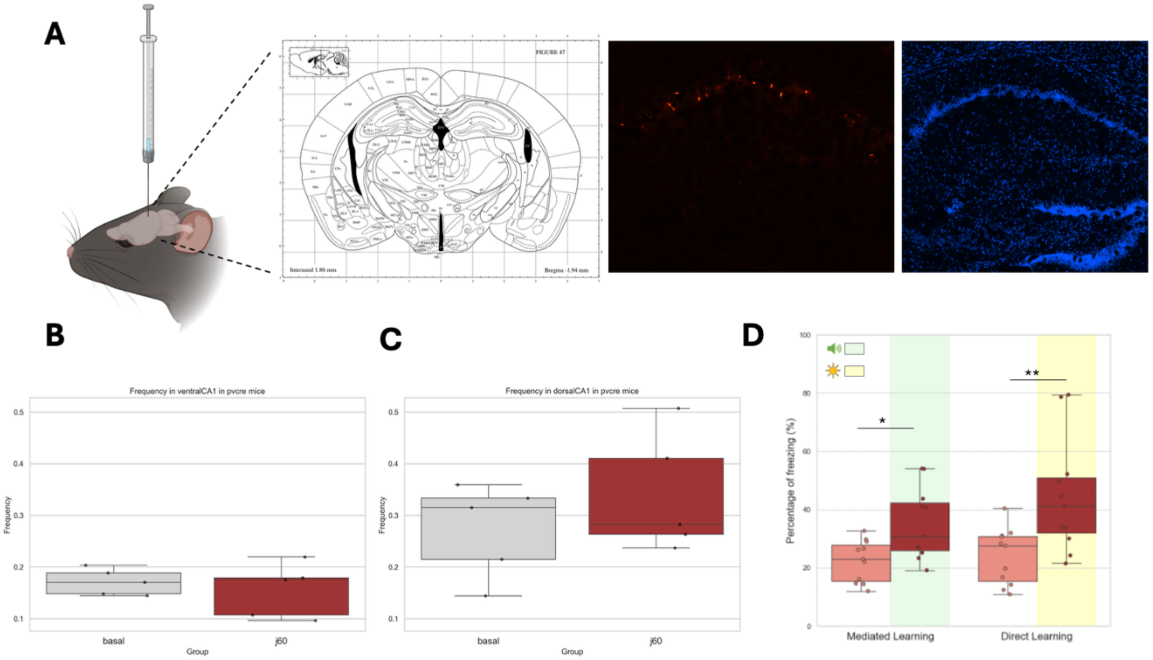


**Supplementary Figure 8 Inhibition of PV-interneurons in dorsal hippocampus during Light-Tone associations did not impact mediated learning.** (A) Schematic representation of the chemogenetic approach in PV-Cre mice, where a cre-dependent inhibitory DREADD was infused in dHPC (representative image on right). Frequency of calcium transients in (B) dHPC and (C) vHPC in PV-positive mice in basal and under J60 conditions. (D) During the preconditioning phase, the DREADD agonist J60 was injected intraperitoneally and the percentage of time spent in freezing during OFF and ON periods of the Probe Test 1 (Mediated learning) and Probe Test 2 (Direct learning) is shown. *, p-value (<0.05); **, p-value (<0.01). Statistical details in Supplementary Table 2.

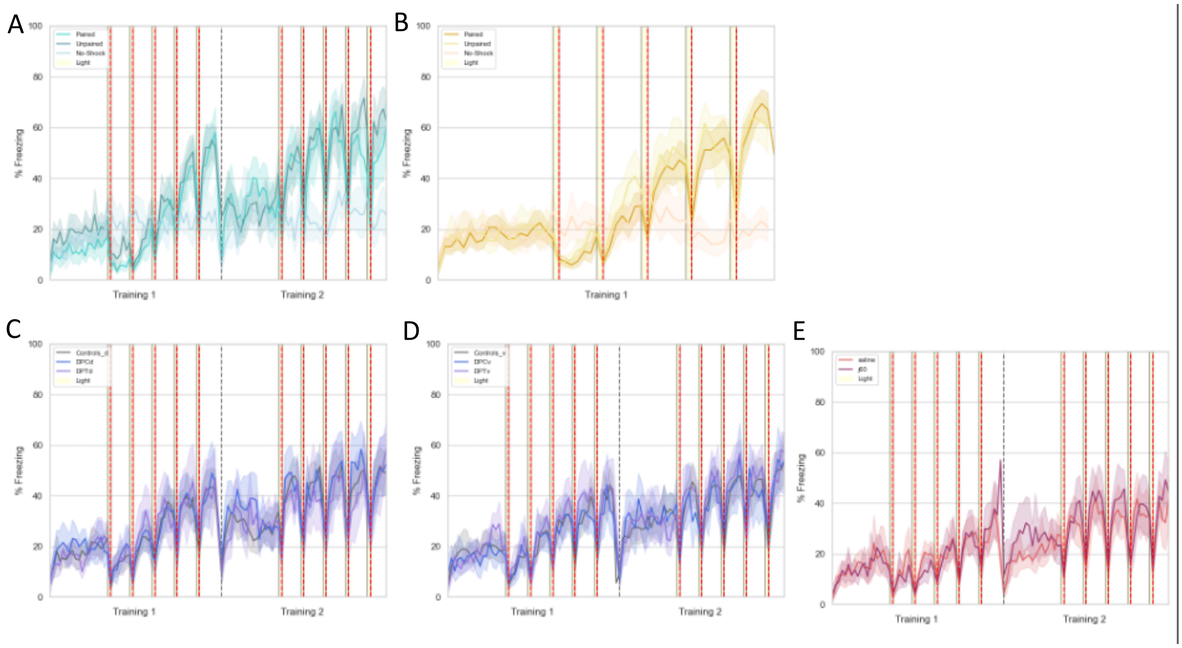


**Supplementary Figure 9. Temporal dynamics of the conditioning phase of all sensory preconditioning experiments.** A) Freezing percentage across the conditioning phase composed by two sessions (Training 1 and Training 2) of the protocol of sensory preconditioning on males with the respective control groups (Paired, Unpaired, and No-Shock). A) Freezing percentage across the conditioning phase composed by one session (Training 1) of the protocol of sensory preconditioning on females with the respective control groups (Paired, Unpaired, and No-Shock). C) Freezing percentage across the conditioning phase composed by two sessions (Training 1 and Training 2) during chemogenetic modulation of dHPC on preconditioning (DPCd) and on probe test (DPTc) compared with controls (Constrols_d) during sensory preconditioning on males. D) Freezing percentage across the conditioning phase composed by two sessions (Training 1 and Training 2) during chemogenetic modulation of vHPC on preconditioning (DPCv) and on probe test (DPTv) compared with controls (Constrols_v) during sensory preconditioning on males. E) Freezing percentage across the conditioning phase composed by two sessions (Training 1 and Training 2) during chemogenetic modulation of CaMKII-positive neurons on dHPC during conditioning phase (J60) compared saline treated animals during sensory preconditioning on males. In yellow bars, the periods of light on, and in red dashed lines, the periods of 2s shock stimulus.

**Table 1.** Statistical analysis. Related to Main Figures 1-4

| Fig. | Condition | n (per group) | Statistical method (interaction main effect) | Factors analysed | Normality and homoscedasticity | Planned Comparisons or Posthoc | P-value |
|---|---|---|---|---|---|---|---|
| 1C | Mediated | 12 | GLM=-5.36, p_value=0.013 | Freezing,experiment(paired, unpaired, no-shock), Off vs On | No | Paired Off vs On | 0,004 |
| | | | | | | Unpaired Off vs On | 0,083 |
| | | | | | | No-shock Off vs On | 0,044 |
| | | | | | | Off Paired vs Unpaired | 1,000 |
| | | | | | | Off Paired vs No-shock | 0,046 |
| | | | | | | Off Unpaired vs No-shock | <0,001 |
| | | | | | | On Paired vs Unpaired | 1,000 |
| | | | | | | On Paired vs No-shock | 0,001 |
| | | | | | | On Unpaired vs No-shock | <0,001 |
| 1E | Direct | 12 | GLM=-18.09, p_value=0.000 | Freezing,experiment(paired, unpaired, no-shock), Off vs On | No | Paired Off vs On | 0,004 |
| | | | | | | Unpaired Off vs On | 0,004 |
| | | | | | | No-shock Off vs On | 0,004 |
| | | | | | | Off Paired vs Unpaired | 0,065 |
| | | | | | | Off Paired vs No-shock | 0,065 |
| | | | | | | Off Unpaired vs No-shock | 0,001 |
| | | | | | | On Paired vs Unpaired | 1,000 |
| | | | | | | On Paired vs No-shock | <0,001 |
| | | | | | | On Unpaired vs No-shock | <0,001 |
| 2B left | Dorsal hippocampus | 7 | Wilcoxon with zero method | Calcium signal (RCaMP or GCaMP), time | No | Gcamp before vs after | 0.0156 |
| | | | | | | Rcamp before vs after | 0.0312 |
| 2C left | Ventral hippocampus | 8 | Wilcoxon with zero method | Calcium signal (RCaMP or GCaMP), time | No | Gcamp before vs after | 0.0156 |
| | | | | | | Rcamp before vs after | 0.0078 |
| 2D | Maximum delta df | Same as 2B and 2C | Wilcoxon with zero method | Calcium signal (RCaMP or GCaMP), time | No | Same as 2B and 2C | |
| 3B | Dorsal hippocampus | 6 | Wilcoxon with zero method | Calcium signal (GCaMP), time | No | Gcamp before vs after | |
| 3C | Ventral hippocampus | 6 | Wilcoxon with zero method | Calcium signal (GCaMP), time | No | Gcamp before vs after | |
| 3D | Maximum delta df | Same as 3B and 3C | Wilcoxon with zero method | Calcium signal (GCaMP), time | No | Same as 3B and 3C | |
| 4B | Mediated | 11-19 | GLM=-1.61, p_value=0.505 | Freezing,dorsal_inhibition(controls, DPC, DPT), Off vs On | No | Controls_d Off vs On | <0,001 |
| | | | | | | DPCd Off vs On | 0,756 |
| | | | | | | DPTd Off vs On | 0,035 |
| | | | | | | Off Controls_d vs DPCd | 0,221 |
| | | | | | | Off Controls_d vs DPTd | 1,000 |
| | | | | | | Off DPCd vs DPTd | 1,000 |
| | | | | | | On Controls_d vs DPCd | 1,000 |
| | | | | | | On Controls_d vs DPTd | 1,000 |
| | | | | | | On DPCd vs DPTd | 1,000 |
| 4C | Mediated | 11-21 | GLM=1.75, p_value=0.539 | Freezing, ventral_inhibition (controls, DPC, DPT), Off vs On | No | Controls_v Off vs On | 0,005 |
| | | | | | | DPCv Off vs On | 0,018 |
| | | | | | | DPTv Off vs On | 0,018 |
| | | | | | | Off Controls_v vs DPCv | 1,000 |
| | | | | | | Off Controls_v vs DPTv | 1,000 |
| | | | | | | Off DPCv vs DPTv | 1,000 |
| | | | | | | On Controls_v vs DPCv | 1,000 |
| | | | | | | On Controls_v vs DPTv | 1,000 |
| | | | | | | On DPCv vs DPTv | 1,000 |

| | | | | | | | |
|---|---|---|---|---|---|---|---|
| 4D | Direct | 11-19 | GLM=3.15, p_value=0.398 | Freezing,dorsal_inhibition(controls, DPC, DPT), Off vs On | No | Controls_d Off vs On | <0,001 |
| | | | | | | DPCd Off vs On | 0,018 |
| | | | | | | DPTd Off vs On | 0,035 |
| | | | | | | Off Controls_d vs DPCd | 1,000 |
| | | | | | | Off Controls_d vs DPTd | 1,000 |
| | | | | | | Off DPCd vs DPTd | 1,000 |
| | | | | | | On Controls_d vs DPCd | 1,000 |
| | | | | | | On Controls_d vs DPTd | 1,000 |
| | | | | | | On DPCd vs DPTd | 1,000 |
| 4E | Direct | 11-21 | GLM=2.46, p_value=0.421 | Freezing, ventral_inhibition (controls, DPC, DPT), Off vs On | No | Controls_v Off vs On | <0,001 |
| | | | | | | DPCv Off vs On | 0,018 |
| | | | | | | DPTv Off vs On | 0,035 |
| | | | | | | Off Controls_v vs DPCv | 1,000 |
| | | | | | | Off Controls_v vs DPTv | 1,000 |
| | | | | | | Off DPCv vs DPTv | 1,000 |
| | | | | | | On Controls_v vs DPCv | 1,000 |
| | | | | | | On Controls_v vs DPTv | 1,000 |
| | | | | | | On DPCv vs DPTv | 1,000 |

**Table 2.** Statistical analysis. Related to Supplementary Figures 1-9

| Fig. | Condition | n (per group) | Analysis | Factors analysed | Normality and homoscedasticity | Comparison | P-value |
|---|---|---|---|---|---|---|---|
| S1 upper | All phases | 12 | Spearman correlation (Eztrack: $R_{males}$=0.91, $R_{females}$=0.86, Manual: $R_{males}$=0.96, $R_{females}$=0.95) | Freezing, Sex, method | No | Deeplacut vs Eztrack males | 0.0001 |
| | | | | | | Deeplacut manual females | 0.0001 |
| S1 bottom | Probe test | 12 | Spearman correlation (Eztrack: $R_{males}$=0.94, $R_{females}$=0.88, Manual: $R_{males}$=0.99, $R_{females}$=0.97) | Freezing, Sex, method | No | Deeplacut vs Eztrack males | 0.0001 |
| | | | | | | Deeplacut manual females | 0.0001 |
| S2C | Mediated | 24 | GLM=-7.06, p_value=0.000 | Freezing,experiment(paired, unpaired, no-shock), Off vs On, females | No | Paired Off vs On | <0,001 |
| | | | | | | Unpaired Off vs On | <0,004 |
| | | | | | | No-shock Off vs On | 1.000 |
| | | | | | | Off Paired vs Unpaired | 1.000 |
| | | | | | | Off Paired vs No-shock | 1.000 |
| | | | | | | Off Unpaired vs No-shock | 1.000 |
| | | | | | | On Paired vs Unpaired | 0.351 |
| | | | | | | On Paired vs No-shock | <0,001 |
| | | | | | | On Unpaired vs No-shock | 1.000 |
| S2E | Direct | 24 | GLM=-14.23, p_value=0.000 | Freezing,experiment(paired, unpaired, no-shock), Off vs On | No | Paired Off vs On | <0,001 |
| | | | | | | Unpaired Off vs On | 0,004 |
| | | | | | | No-shock Off vs On | 0,242 |
| | | | | | | Off Paired vs Unpaired | 1,000 |
| | | | | | | Off Paired vs No-shock | 1,000 |
| | | | | | | Off Unpaired vs No-shock | 0,545 |
| | | | | | | On Paired vs Unpaired | 1,000 |
| | | | | | | On Paired vs No-shock | <0,001 |
| | | | | | | On Unpaired vs No-shock | 0,001 |
| S3A | Mediated, Direct | 12 | Spearman correlation (Mediated vs Direct: $R_{males}$=0.70) | Freezing, Sex | No | 0.011 | |
| S3B | Mediated, Direct | 24 | Spearman correlation (Mediated vs Direct: $R_{females}$=0.02) | Freezing, Sex | No | 0.920 | |
| S4B left | Dorsal hippocampus | 7 | Wilcoxon with zero method | Calcium signal (Rcamp or Gcamp), time | No | Gcamp before vs after | 0.015625 |
| | | | | | | Rcamp before vs after | 0.03125 |
| S4C left | Ventral hippocampus | 8 | Wilcoxon with zero method | Calcium signal (Rcamp or Gcamp), time | No | Gcamp before vs after | 0.015625 |
| | | | | | | Rcamp before vs after | 0.0234375 |

| | | | | | | | |
|---|---|---|---|---|---|---|---|
| S4D | Maximum delta df | Same as S4B and C | Wilcoxon with zero method | Calcium signal (Rcamp or Gcamp), time, dorsal and ventral hippocampus | No | Same as S4B and C | |
| S5B | Dorsal hippocampus | 7 | Wilcoxon with zero method | Calcium signal (Rcamp or Gcamp), time | No | Gcamp before vs after | 0.015625 |
| | | | | | | Rcamp before vs after | 0.03125 |
| S5C | Ventral hippocampus | 8 | Wilcoxon with zero method | Calcium signal (Rcamp or Gcamp), time | No | Gcamp before vs after | 0.015625 |
| | | | | | | Rcamp before vs after | 0.0234375 |
| S5D | Maximum delta df | Same as S3B and C | Wilcoxon with zero method | Calcium signal (Rcamp or Gcamp), time, dorsal and ventral hippocampus | No | Same as S3B and C | |
| S6B | Dorsal hippocampus | 7 | Wilcoxon with zero method | Calcium signal (Rcamp or Gcamp), time | No | Gcamp before vs after | 0.015625 |
| | | | | | | Rcamp before vs after | 0.03125 |
| S6C | Ventral hippocampus | 8 | Wilcoxon with zero method | Calcium signal (Rcamp or Gcamp), time | No | Gcamp before vs after | 0.015625 |
| | | | | | | Rcamp before vs after | 0.0078125 |
| S6D | Maximum delta df | Same as S5B and C | Wilcoxon with zero method | Calcium signal (Rcamp or Gcamp), time, dorsal and ventral hippocampus | No | Same as S4B and C | |
| S7A | Mediated | 21 | Kruskall-Walis=4.73 | Freezing, dorsal controls groups (CS, CJ60, DS) | No | 0.094 | |
| S7B | Direct | 21 | Kruskall-Walis=0.99 | Freezing, dorsal controls groups (CS, CJ60, DS) | No | 0.610 | |
| S7C | Mediated | 19 | Kruskall-Walis=0.58 | Freezing, ventral controls groups (CS, CJ60, DS) | No | 0.748 | |
| S7D | Direct | 19 | Kruskall-Walis=1.99 | Freezing, ventral controls groups (CS, CJ60, DS) | No | 0.370 | |
| S7E | Calcium frequency | 4 | Wilcoxon test | Basal vs j60 | No | 0.25 | |
| S7F | Calcium frequency | 4 | Wilcoxon test | Basal vs j60 | No | 0.25 | |
| S7G | Freezing | 22 | Mixed-effects analysis followed by Tuke's multiple comparisons test | Freezing and drug | No | Saline : Off vs On | 0.0006 |
| | | | | | No | J60: Off vs On | 0.012 |
| | | | | | No | On : saline vs J60 | 0.024 |
| S8B | Calcium frequency | 4 | Wilcoxon test | Basal vs j60 | No | 0.187 | |
| S8C | Calcium frequency | 4 | Wilcoxon test | Basal vs j60 | No | 0.625 | |
| S8D | Mediated | 11 | GLM= -16.57, p value=0.001 | Freezing | No | Off vs On | 0.049 |
| S8D | Direct | 11 | GLM= -16.57, p value=0.001 | Freezing | No | Off vs On | 0.00 |
